# Light-sheet excitation-encoded volumetric spectroscopy for fast multiplexed imaging and quantitative physicochemical mapping in cells

**DOI:** 10.64898/2026.04.14.718435

**Authors:** Jinhong Yan, Jiankai Xia, Jiayi Liu, Yi He, Jeongmin Kim, Kun Chen

## Abstract

Understanding how cellular organization relates to function requires imaging approaches that resolve structural and physicochemical information throughout three-dimensional volumes. Here we introduce light-sheet excitation spectroscopic microscopy (LS-ExSM), a method that captures full excitation spectra with ∼10 nm spectral resolution at each voxel for multiplexed imaging and quantitative spectroscopic readout. Excitation-wavelength encoding makes this spectroscopic readout compatible with single-objective wide-field light-sheet imaging while maintaining high spectral fidelity. Combined with sparse volumetric sampling and deep-learning-based reconstruction, LS-ExSM achieves high-speed volumetric imaging while preserving structural and spectral information. We demonstrate six-color volumetric imaging of live cells at near-second temporal resolution with minimal crosstalk, enabling visualization and quantitative analysis of dynamic organelle interactions in three dimensions. We further apply LS-ExSM to three-dimensional mapping of lipid-droplet polarity at near-diffraction-limited resolution. Together, these results establish excitation-encoded volumetric spectroscopy as a route to whole-cell imaging of subcellular organization and physicochemical state.

## Introduction

A comprehensive understanding of cellular organization and function requires imaging approaches that capture biological processes within their native three-dimensional (3D) context^1^. Beyond resolving cellular architecture, 3D imaging is also needed to map the spatial organization of molecular and physicochemical states that underlie functional heterogeneity across cellular volumes^2, 3^. By selectively illuminating only the focal plane, light-sheet microscopy has emerged as a powerful platform for this purpose because it combines high speed, efficient photon usage and reduced phototoxicity^4–9^. Recent advances in single-objective and oblique-plane implementations have further improved experimental accessibility and compatibility with conventional sample preparations ^10–13^, facilitating routine whole-cell imaging^14–19^. However, most current light-sheet methods remain limited in multiplexing capacity, typically visualizing only two or three structures simultaneously. Because cellular function arises from coordinated interactions among multiple molecular species and organelles, there is a growing need for imaging strategies that support multiplexed visualization within the same 3D volume^20^. Fluorescence spectral imaging offers a scalable route toward this goal by exploiting spectral differences among labels^21, 22^, while also enabling quantitative readout of local physicochemical states using environmentally responsive probes^23^.

In light-sheet microscopy, emission-based spectral imaging is fundamentally constrained by the difficulty of acquiring spatial and spectral information simultaneously in a wide-field detection geometry. Spectrally resolved imaging therefore typically relies on point- or line-scanning approaches based on fluorescence dispersion^20, 24^, which constrain volumetric imaging speed. Recent parallel detection schemes enable faster multispectral imaging by distributing fluorescence across multiple channels, but generally provide only coarse spectral sampling (>30 nm), reducing spectral fidelity and signal-to-noise ratio^25^. As a result, high-resolution spectral measurements—and thus quantitative spectroscopic interrogation in three dimensions—remain challenging. Excitation-based spectral microscopy addresses these limitations by encoding spectral information through rapid modulation of excitation wavelengths while collecting emission within a fixed passband^26, 27^. This strategy preserves photon efficiency, avoids dependence on the spectral response of the detection system and is inherently compatible with wide-field light-sheet imaging. Previous excitation-based implementations combined with lattice light-sheet microscopy enabled faster volumetric imaging, but relied on discrete excitation wavelengths and wavelength-dependent illumination profiles, limiting spectral resolution and increasing crosstalk among cellular structures^20^. More recently, excitation spectral microscopy (ExSM) enabled multiplexed imaging with minimal target-to-target crosstalk together with sensitive quantification of local physicochemical states from high-fidelity excitation spectra^26, 28, 29^. However, extending this capability to volumetric light-sheet imaging while preserving spectral resolution, multiplexing capacity and imaging speed across whole cells has remained an unresolved challenge.

Here we introduce light-sheet excitation spectroscopic microscopy (LS-ExSM), an excitation-encoded approach for volumetric spectroscopic imaging in live cells. By capturing full excitation spectra with ∼10 nm spectral resolution at each voxel, LS-ExSM enables multiplexed imaging together with quantitative mapping of physicochemical states across three-dimensional cellular volumes. Excitation-wavelength encoding makes this spectroscopic readout compatible with single-objective wide-field light-sheet imaging while maintaining high spectral fidelity, and sparse volumetric sampling combined with deep-learning-based reconstruction further enables rapid volumetric imaging. We apply LS-ExSM to fast six-color live-cell imaging with minimal crosstalk and to quantitative mapping of lipid droplet (LD) polarity in three dimensions. These results establish excitation-encoded light-sheet microscopy as a general strategy for linking three-dimensional cellular organization to local physicochemical states.

## Results

### LS-ExSM enables multiplexed volumetric imaging of subcellular structures

We developed a volumetric excitation–spectral imaging platform by integrating excitation spectral microscopy with single-objective light-sheet illumination (LS-ExSM). In this system, excitation wavelengths from a broadband supercontinuum source were rapidly tuned using an acousto-optic tunable filter (AOTF) to encode spectral information^28, 29^. In parallel, single-objective light-sheet illumination was implemented in an open-top oblique-plane configuration with a remote imaging module, enabling efficient fluorescence collection with a high-numerical-aperture objective and near-diffraction-limited volumetric imaging in standard sample geometries (Supplementary Fig. S1)^13^. This design decouples spectral encoding from detection, enabling wide-field volumetric imaging with near-diffraction-limited resolution in all three dimensions (*x*: 3249 ± 30 nm; *y*: 328 ± 28 nm; *z*: 509 ± 52 nm after deconvolution), with effective detection numerical apertures of 1.20 along *x* and 1.35 along *y* (Supplementary Figs. S2–S3), while capturing high-fidelity excitation spectra with ∼10 nm spectral resolution at each voxel.

On this platform, spectral and volumetric information were acquired in a unified manner (Fig. 1a). At each light-sheet position, excitation-resolved image stacks were recorded, and the light sheet was then scanned through the sample over approximately 100 planes at 0.373 μm intervals to generate a volumetric spectral dataset. These data formed a four-dimensional excitation-resolved volume, which could be further extended to five dimensions (x–y–z–λ–t) through repeated acquisition over time (see Methods). This data structure enabled voxel-wise analysis of excitation spectra for resolving multiple subcellular structures, while the same spectral information provided quantitative functional readout, thereby establishing LS-ExSM as a spectroscopic framework for volumetric imaging.

**Fig. 1.**
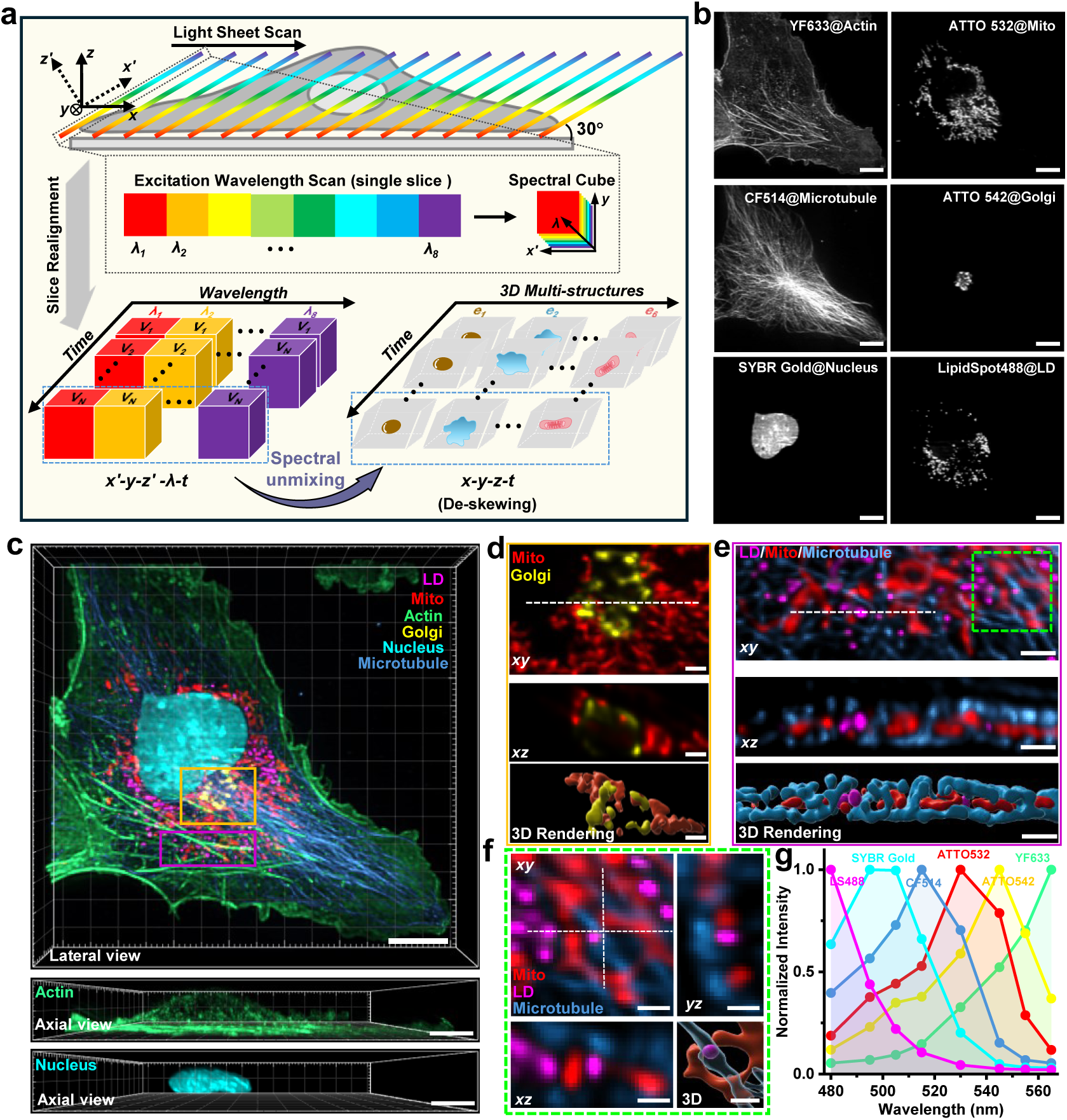
| Oblique single-objective light-sheet illumination combined with excitation spectral microscopy enables multiplexed volumetric imaging of subcellular structures. (a) Schematic of excitation–spectral light-sheet acquisition. A tilted light sheet is scanned across the specimen along the x direction using a galvo mirror, while excitation-resolved image stacks are acquired at each position and assembled into a volumetric spectral dataset. After realignment, this yields a four-dimensional excitation-resolved volume (x′–y–z′–λ) at each time point, extendable to five dimensions (x′–y–z′–λ–t) through repeated acquisition. Voxel-wise spectral unmixing separates individual structures, and de-skewing produces geometrically corrected three-dimensional images (x–y–z). (b) Spectrally unmixed images (maximum intensity projections) of six subcellular structures in a fixed COS-7 cell labeled with spectrally overlapping fluorophores. (c) Three-dimensional rendering of the unmixed structures, showing a lateral view of the whole cell and axial views of actin and nucleus. (d) Magnified region from (c) (yellow box) showing mitochondria–Golgi organization. Orthogonal xy and xz sections (along the indicated dashed line) and surface rendering reveal their partially overlapped yet distinct three-dimensional arrangement. (e) Enlarged view from (c) (magenta box) showing mitochondria and LDs embedded within the microtubule network, with corresponding orthogonal sections and 3D surface rendering. (f) Zoomed-in view from (e) highlighting mitochondria–LD contact sites. Orthogonal xy, xz and yz sections together with surface rendering reveal their three-dimensional proximity. (g) High-fidelity excitation spectra of the six fluorophores, separately measured on our setup using singly labeled samples. Scale bars, 10 μm (b, c), 2 μm (d, e), 1 μm (f).

We next evaluated multiplexed volumetric imaging in fixed COS-7 cells labeled with six spectrally overlapping fluorophores targeting major subcellular structures, including actin, microtubules, mitochondria, Golgi apparatus, nucleus and LDs (Fig. 1b–g). Spectral unmixing based on reference excitation spectra enabled clear separation of all six structures with minimal target-to-target crosstalk averaging less than 1% (Fig. 1b and Supplementary Fig. S4), consistent with reduced ambiguity from axial overlap in volumetric acquisition. Three-dimensional rendering of the unmixed signals revealed the global organization of the cell with subcellular resolution (Fig. 1c and Supplementary Video 1), enabling reliable visualization of fine structural features throughout the cellular volume. Cross-sectional views further highlighted diverse spatial relationships among organelles in three dimensions (Supplementary Fig. S5).

Volumetric imaging further enabled direct interrogation of spatial relationships that are difficult to resolve in two-dimensional projections. Mitochondria and Golgi showed partially overlapping yet distinct distributions in orthogonal sections and three-dimensional renderings (Fig. 1d). Similarly, mitochondria and LDs were embedded within the microtubule network, forming complex spatial arrangements that are not apparent in planar views (Fig. 1e). At mitochondria–LD contact sites, LDs aligned along microtubules while mitochondria partially wrapped around them, forming a grasping-like geometry resolved in three dimensions (Fig. 1f). Consistent results were observed across additional cells (Supplementary Fig. S6). Together, these results establish LS-ExSM as a robust platform for multiplexed spectroscopic volumetric imaging, enabling accurate three-dimensional mapping of subcellular organization and interactions. However, achieving this capability currently requires dense sampling across both spatial and spectral dimensions, which limits imaging speed and motivates strategies to reduce acquisition redundancy while preserving volumetric and spectral information.

### Sparse volumetric sampling with deep-learning reconstruction accelerates LS-ExSM for high-speed spectroscopic imaging

To overcome the speed limitation imposed by dense sampling across spatial and spectral dimensions, we developed a sparse acquisition strategy coupled with deep learning–based reconstruction (Fig. 2a–b). Rather than imaging every light-sheet position, only a subset of slices was acquired at larger intervals (for example, every third plane), increasing volumetric imaging speed by threefold. Missing information was recovered using a novel two-stage neural network framework (Fig. 2b, Supplementary Figs. S7–S8 and Supplementary Note 1–2), comprising a self-supervised denoising network to enhance signal quality, followed by a supervised reconstruction network with edge-aware constraints to infer intermediate planes and restore complete volumes.

**Fig. 2.**
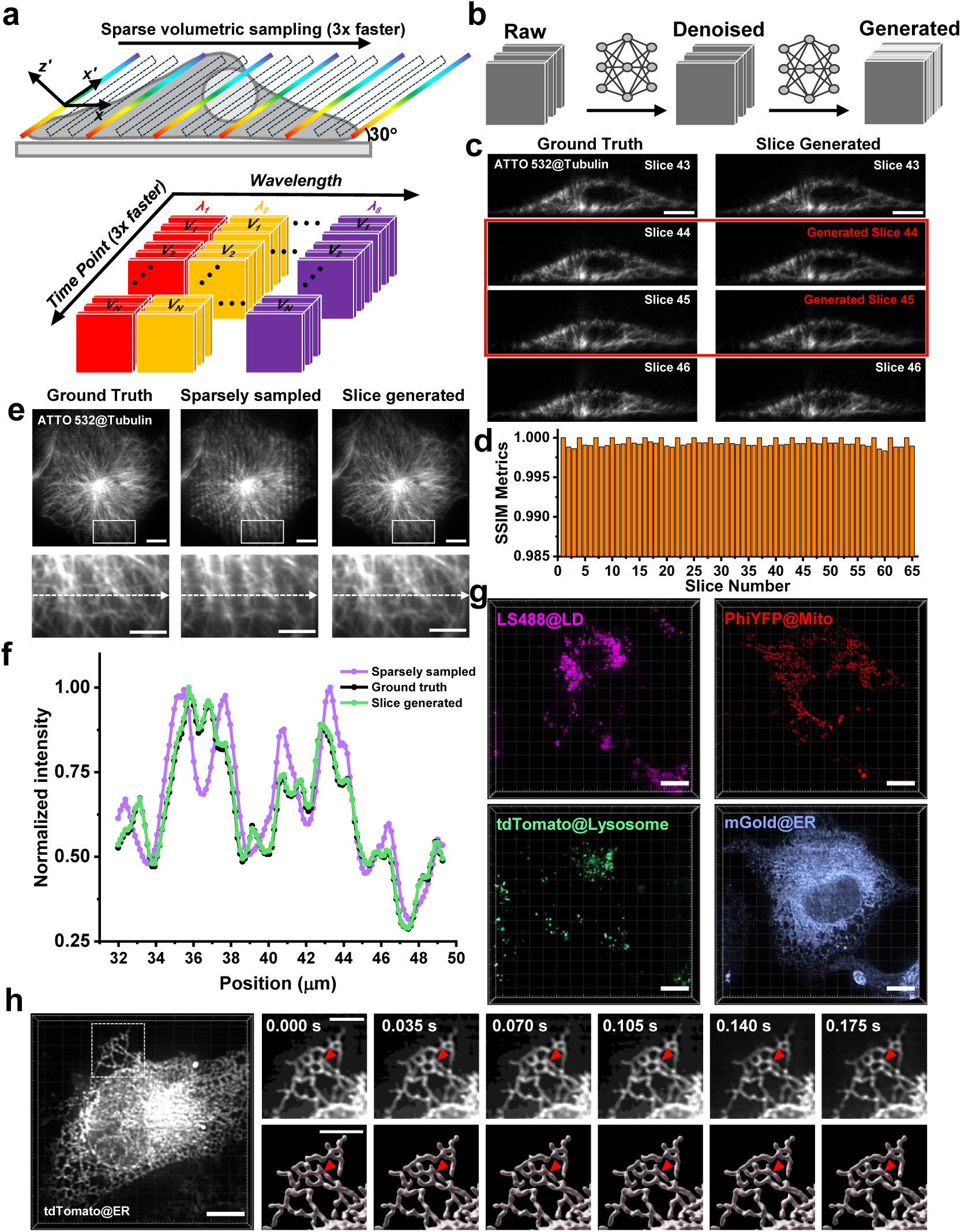
**| Sparse volumetric sampling with deep learning–based reconstruction enables high-speed LS-ExSM while preserving structural and spectral fidelity.** (a) Schematic of sparse light-sheet acquisition. A subset of light-sheet positions is sampled at regular intervals, while omitted planes (dashed boxes) correspond to positions in dense sampling. The resulting five-dimensional dataset (x′–y–z′–λ–t) contains fewer slices per volume, enabling an approximately threefold increase in imaging speed. (b) Reconstruction framework. A self-supervised denoising network enhances signal quality in sparsely sampled data, followed by a supervised reconstruction network that infers missing slices to recover complete volumetric datasets. (c) Comparison between ground truth and reconstructed slices in a fixed COS-7 cell labeled for microtubules. Reconstructed slices are indicated by red boxes. (d) Quantitative comparison of generated and ground-truth slices using the structural similarity index (SSIM), showing consistently high similarity (SSIM > 0.998) across all evaluated slices. (e) Maximum intensity projections of microtubules for ground truth, sparse input and reconstructed volumes. Insets (white boxes) highlight recovery of fine structures lost in sparse sampling. (f) Line profiles along the indicated dashed line in (e), showing close agreement between reconstructed and ground-truth signals and loss of detail in the sparse input. (g) Spectrally unmixed images of four subcellular structures in a live COS-7 cell, reconstructed from sparsely sampled volumetric data, demonstrating preserved spectral separation and three-dimensional organization. (h) High-speed volumetric imaging of the endoplasmic reticulum (ER) network in live COS-7 cells. Time-lapse views of the boxed region (white dashed lines) reveal rapid three-dimensional remodeling enabled by accelerated acquisition. Scale bars, 10 μm (c, e top, g, h left), 5 μm (h right, e bottom).

We next evaluated reconstruction accuracy using fixed cells labeled with a microtubule marker (Fig. 2c–f). Fully sampled datasets were used as ground truth, from which sparsely sampled inputs were generated for reconstruction. The reconstructed slices closely matched the ground truth both visually and quantitatively (Fig. 2c–d), with structural similarity index (SSIM) values consistently exceeding 0.998, indicating high structural fidelity. Line profiles extracted from representative cross-sections further confirmed that structural features were preserved, whereas sparsely sampled data alone showed substantial information loss (Fig. 2e–f). We then assessed whether spectral information was preserved during reconstruction. Excitation spectra extracted from corresponding voxels in reconstructed and ground-truth volumes showed excellent agreement across all wavelengths (Supplementary Fig. S9), with pixel-wise spectral angle mapping (SAM) values predominantly below 1°, indicating that high-fidelity excitation spectra were preserved after reconstruction.

Consistent with this, four-color live-cell imaging demonstrated reliable spectral unmixing with an average target-to-target crosstalk below 1% while preserving three-dimensional organization and dynamics (Fig. 2g, Supplementary Fig. S10 and Supplementary Video 2), indicating that sparse sampling did not compromise spectral fidelity. This robustness was further supported by the denoising capability of the deep-learning reconstruction network, which improved the signal-to-noise ratio of raw slices by ∼10 dB (Supplementary Fig. S11). Within this framework, LS-ExSM achieved ultrafast volumetric imaging of the endoplasmic reticulum (ER) at 28 Hz (35 ms per volume; ∼1 ms exposure time for each optical section) (Fig. 2h and Supplementary Video 3), enabling direct visualization of rapid three-dimensional remodeling of ER tubules.

### High-speed volumetric LS-ExSM enables 6-color imaging of dynamic subcellular interactions in live cells

Building on the accelerated framework, we performed six-color volumetric imaging of live COS-7 cells labeled with spectrally overlapping probes targeting major organelles, including lysosomes, peroxisomes, mitochondria, ER, DNA and LDs (Fig. 3 and Supplementary Video 4). Volumetric imaging was achieved at near-second temporal resolution (∼1.2 s per volume) over 200 consecutive time points. This temporal resolution was primarily selected to balance signal-to-noise ratio and phototoxicity in live cells (see Methods), with potential for further acceleration using brighter probes.

**Fig. 3.**
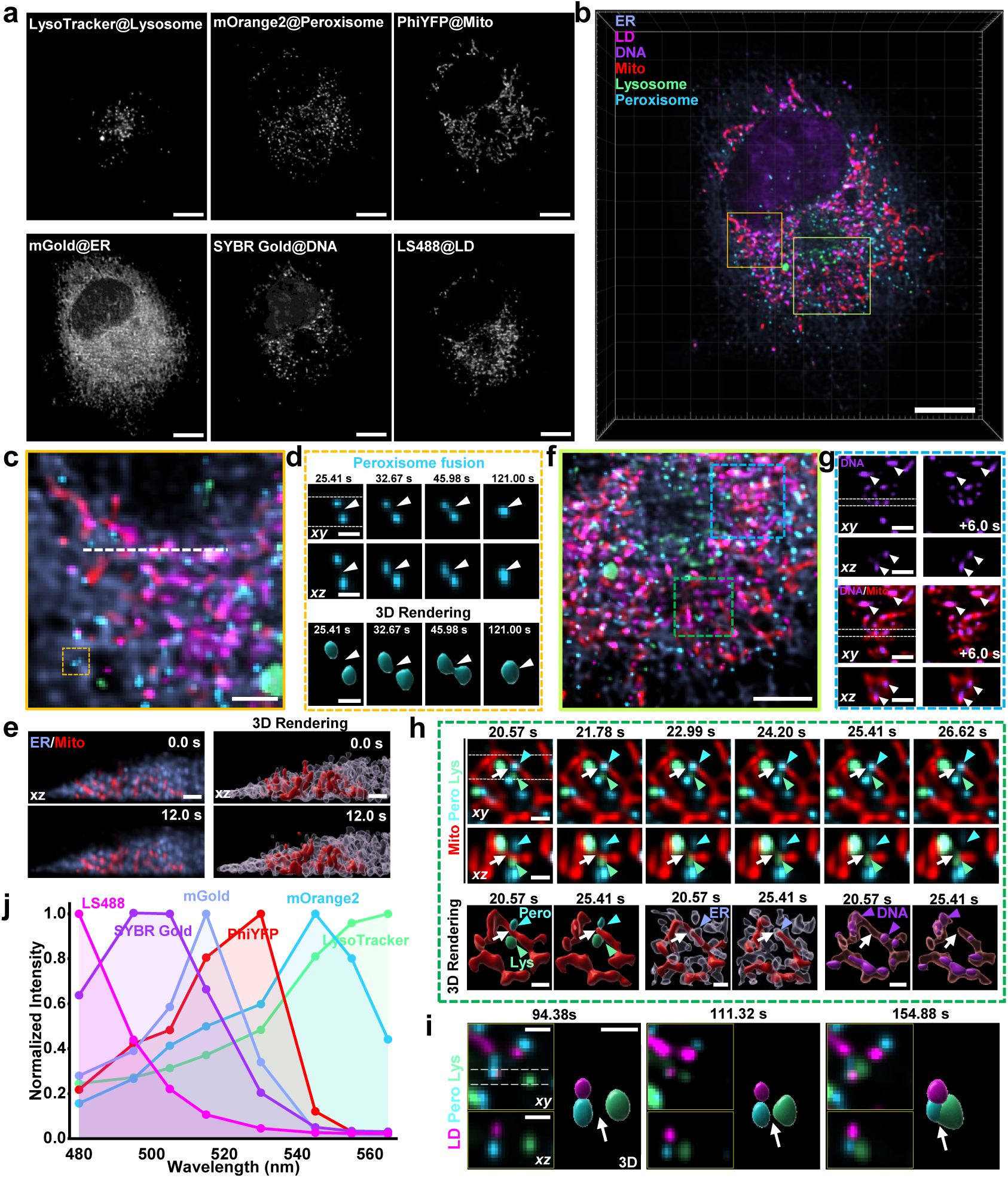
**| High-speed LS-ExSM enables multiplexed imaging of dynamic multi-organelle interactions in live cells.** (a) Spectrally unmixed images of six subcellular structures in a live COS-7 cell labeled with spectrally overlapping fluorophores. (b) Three-dimensional maximum intensity projection of the six structures, showing their spatial organization within the same cellular volume. Colored boxes indicate regions selected for detailed analysis in (c-i). (c) Magnified view of a selected region showing peroxisome dynamics. (d) Further enlargement of the dashed region in (c), showing a peroxisome fusion event. Orthogonal xy and xz sections and corresponding three-dimensional renderings at representative time points illustrate progressive merging of the two organelles. (e) Cross-sectional view along the indicated dashed line showing mitochondria and endoplasmic reticulum (ER) organization. Orthogonal sections and three-dimensional renderings reveal close spatial association, with mitochondria partially enveloped by ER tubules. (f) Magnified view of the region indicated in (b) (green box). (g) Enlargement of the dashed region in (f), showing mtDNA localized within mitochondria. Orthogonal xy and xz sections at two time points illustrate persistent co-localization. (h) Time-lapse imaging of a mitochondrial fission event from the region indicated in (f) (green dashed box). Orthogonal views and three-dimensional renderings show localization of lysosomes and peroxisomes near the constriction site, with ER tubules intersecting the same region. (i) Time-lapse visualization of interactions among LDs, peroxisomes and lysosomes, showing the formation of multi-organelle assemblies. (j) High-fidelity excitation spectra of the six fluorophores used for spectral unmixing. Scale bars, 10 μm (a, b), 5 μm (f), 2 μm (c, e, g), 1 μm (d, h, i).

The multiplexed volumetric data enabled direct visualization of dynamic interactions and spatial organization among multiple organelles within the same cellular volume. For example, rapid fusion of peroxisomes was clearly resolved in orthogonal sections and three-dimensional renderings (Fig. 3c–d and Supplementary Video 5). Cross-sectional views also revealed extensive spatial coupling between mitochondria and the ER network, with mitochondria closely associated with ER tubules throughout the cellular volume (Fig. 3e). The high spatiotemporal resolution further enabled tracking of coordinated dynamics across multiple organelles. Co-localization of mitochondrial DNA (mtDNA) within mitochondria could be resolved in three dimensions and followed over time (Fig. 3f–g), supporting consistent labeling during dynamic processes. Mitochondrial fission events were directly visualized, in which lysosomes and peroxisomes localized to opposite sides of mitochondrial constriction sites, while ER tubules intersected the same regions and mtDNA was partitioned into daughter mitochondria (Fig. 3h). These observations were consistent with previous reports of ER- and lysosome-associated mitochondrial division^30–33^ and suggested a potential involvement of peroxisomes in this process, beyond their established role in sharing fission machinery with mitochondria^34–36^. Multiplexed volumetric imaging also revealed closely associated multi-organelle assemblies involving LDs, peroxisomes and lysosomes (Fig. 3i and Supplementary Video 6), illustrating the ability of LS-ExSM to resolve complex spatial organization in live cells. The spectral identities of all labeled structures were well defined by their excitation profiles (Fig. 3j), enabling robust separation of multiple targets throughout the volumetric time series. Together, these results demonstrate that high-speed LS-ExSM enables simultaneous three-dimensional visualization of interacting organelles in live cells, providing access to dynamic processes that are difficult to capture with conventional approaches.

### LS-ExSM reveals three-dimensional organelle interactions under metabolic perturbations

Organelle interactions are dynamically regulated by cellular metabolic states, yet their organization in three dimensions remains incompletely understood. To address this, we used LS-ExSM to map contacts among LDs, lysosomes and mitochondria under control, starvation and ATGL inhibition^37^ (Atglistatin-treated, ATGLi+) conditions. Volumetric datasets were surface-segmented, and pairwise organelle proximity was quantified in three dimensions (Fig. 4a, see Methods). Notably, contacts inferred from two-dimensional projections were not always preserved in three dimensions, whereas true spatial contacts remained consistent across viewing angles.

**Fig. 4.**
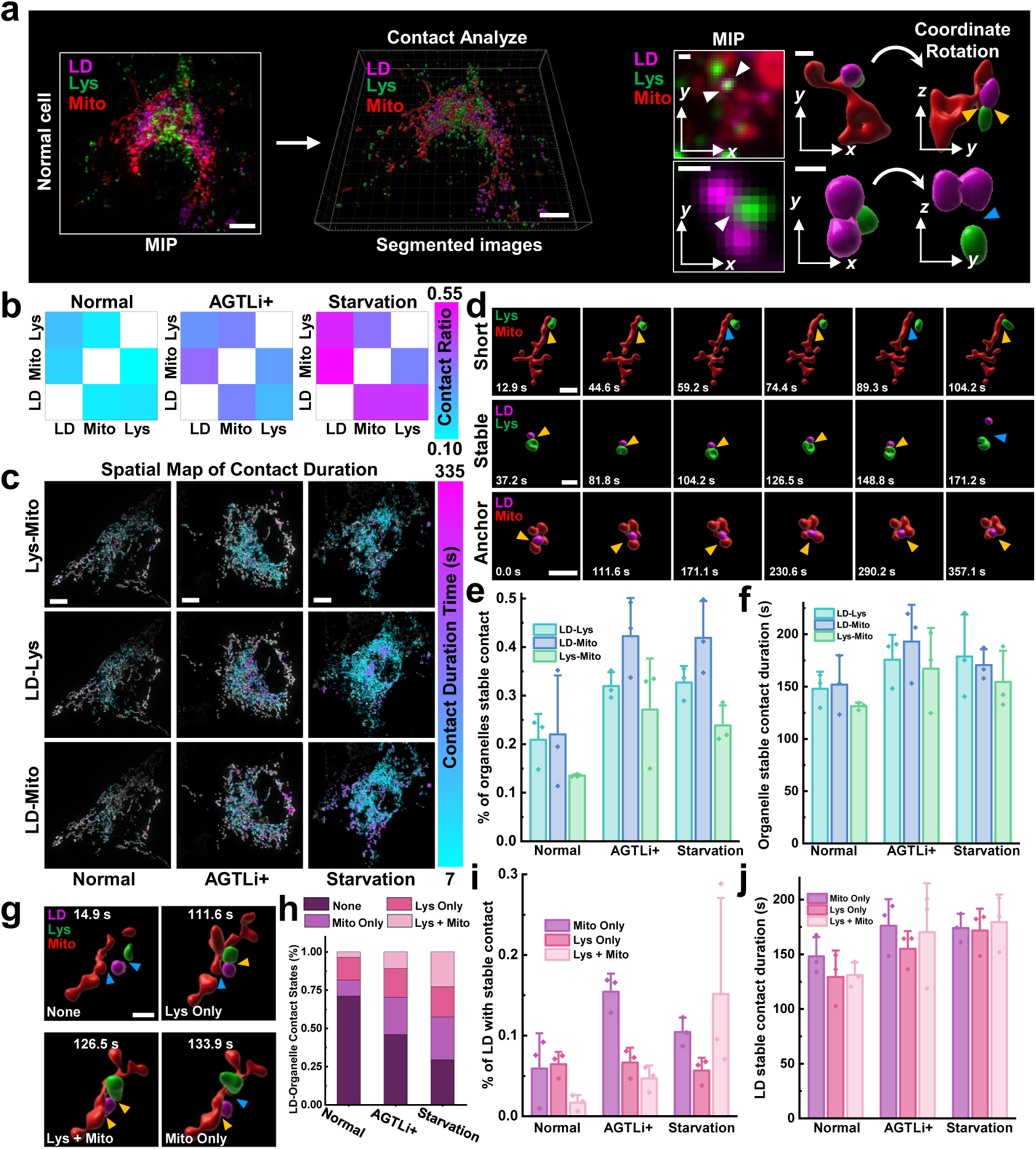
**| LS-ExSM enables quantitative analysis of three-dimensional multi-organelle contact dynamics under metabolic perturbations.** Workflow for three-dimensional contact analysis of LDs labelled with LS488, mitochondria labelled with PhiYFP, and lysosomes labelled with tdTomato. Organelle surfaces are segmented from volumetric data to quantify pairwise proximity. Comparison of maximum intensity projection and rotated three-dimensional views shows that apparent contacts in two-dimensional projections may not correspond to true spatial contacts. (b) Mean per-frame contact ratios for organelle pairs (LD–mitochondria, LD–lysosome and mitochondria–lysosome) under normal, ATGLi+ and starvation conditions. (c) Spatial maps of contact sites projected in two dimensions, with color indicating contact duration for individual interaction events. (d) Representative examples of contact categories defined by duration: short (<60 s), stable (>60 s) and anchored contacts (persisting throughout the observation window). (e) Fraction of organelles exhibiting at least one stable contact (≥60 s) for each organelle pair across conditions. (f) Mean total duration of stable contacts (>60 s) per organelle, averaged over organelles with at least one such contact, for each organelle pair across conditions. (g) Representative LD contact states, including no contact, lysosome-only, mitochondria-only and simultaneous contacts with both organelles. (h) Mean per-cell LD state occupancy, calculated as the fraction of time each LD spends in each contact state and averaged across LDs within each cell, under different metabolic conditions. (i) Fraction of LDs exhibiting at least one stable contact (≥60 s) in each contact state (mitochondria-only, lysosome-only and dual contacts) across conditions. (j) Mean total duration of stable contacts (>60 s) per LD in each contact state, averaged over LDs with at least one such contact, across conditions. n = 3 cells. Error bars indicate standard error of the mean (s.e.m.). Scale bars, 10 μm (a left, c), 2 μm (d), 1 μm (g), 0.5 μm (a right).

We first quantified the frequency of pairwise contacts across conditions based on volumetric analysis (Fig. 4b and Supplementary Fig. S12). Both starvation and ATGL inhibition increased overall contact frequency, indicating enhanced organelle proximity under metabolic perturbation. This increase was most pronounced for LD interactions with mitochondria and lysosomes during starvation, consistent with increased coordination among lipid storage, energy production and degradation pathways. Importantly, quantitative comparison further showed that two-dimensional analysis substantially overestimated contact frequency and could even reverse biological trends—for example, LD–mitochondria contacts appeared reduced under starvation in 2D but increased in 3D (Supplementary Fig. S13)—highlighting the necessity of volumetric measurements for accurate quantification of organelle contacts. Spatial maps further revealed that contacts were broadly distributed throughout the cellular volume but exhibited substantial heterogeneity in both frequency and duration (Fig. 4c). To further characterize interaction dynamics, contacts were classified by duration into short (<60 s) and stable (>60 s) events^32, 38, 39^, with persistent contacts spanning the entire observation window defined as anchored (Fig. 4d). Short contacts likely reflected transient encounters, whereas stable contacts indicated more sustained interactions. Both starvation and ATGL inhibition increased not only overall contact frequency but also the fraction and duration of stable contacts, with the strongest effect observed for LD–mitochondria interactions (Fig. 4e–f). Increased stability and duration under starvation were consistent with enhanced metabolic coupling, likely reflecting elevated demand for lipid mobilization. By contrast, although ATGL inhibition blocks neutral lipolysis, it still promoted stable and prolonged contacts, suggesting that while organelles were efficiently recruited into contact, the absence of lipid hydrolysis may delay their resolution.

We next examined how these interactions were organized around individual LDs. LDs dynamically transitioned among four contact states: no contact, lysosome-only, mitochondria-only, and simultaneous contacts with both organelles (Fig. 4g). Under both starvation and ATGL inhibition, the fraction of LDs engaging mitochondria—either alone or together with lysosomes—increased markedly, accompanied by a reduction in the no-contact state (Fig. 4h), with the strongest shift observed under starvation. Analysis of stable interactions revealed distinct LD-centered reorganization patterns (Fig. 4i). Under ATGL inhibition, the increase was dominated by mitochondria-only stable contacts, whereas lysosome-only contacts remained largely unchanged. In contrast, starvation preferentially increased stable lysosome–mitochondria co-associated contacts, with a smaller increase in mitochondria-only interactions. Contact durations were also prolonged under both conditions (Fig. 4j). These patterns were consistent with enhanced lysosome–mitochondria coordination around LDs under starvation, whereas under ATGL inhibition, persistent mitochondria-associated contacts may reflect delayed progression of lipid-processing steps when neutral lipolysis is blocked.

### Three-dimensional quantitative mapping of LD polarity

LD polarity has previously been established as a sensitive reporter of cellular metabolic state^29^. Here, we extended LS-ExSM to enable volumetric mapping of LD polarity using the solvatochromic dye Nile Red, providing direct spectroscopic readout of physicochemical states across entire cellular volumes. This capability arises from capturing high-fidelity excitation spectra with ∼10 nm spectral resolution and remains largely inaccessible with existing volumetric imaging approaches. To establish ground truth, we constructed model systems consisting of pure triolein droplets and artificial adiposomes with homogeneous lipid cores (triolein) and defined phospholipid monolayers (DOPC; 1,2-dioleoyl-sn-glycero-3-phosphocholine). Three-dimensional excitation spectroscopic imaging revealed subtle spectral differences between droplet interiors and surfaces. In adiposomes, the surface exhibited an excitation spectrum red-shifted by ∼10 nm relative to the core, consistent with the higher polarity of the DOPC monolayer, although the measured surface signal represents a diffraction-limited mixture of DOPC periphery and triolein core, whereas pure triolein droplets remained spectrally uniform (Fig. 5a). Spectral phasor analysis was used to convert excitation spectra into a single quantitative parameter: the phase angle, which scales linearly with the mean excitation wavelength and reports the local polarity of the dye environment (Fig. 5b). Mapping this parameter across voxels enabled direct visualization of polarity distributions in three dimensions. Triolein droplets exhibited uniform polarity, whereas adiposomes showed a pronounced core–surface gradient (Fig. 5c, Supplementary Fig. S14 and Supplementary Video 7), demonstrating that LS-ExSM resolved physicochemical heterogeneity within individual droplets in three dimensions.

**Fig. 5.**
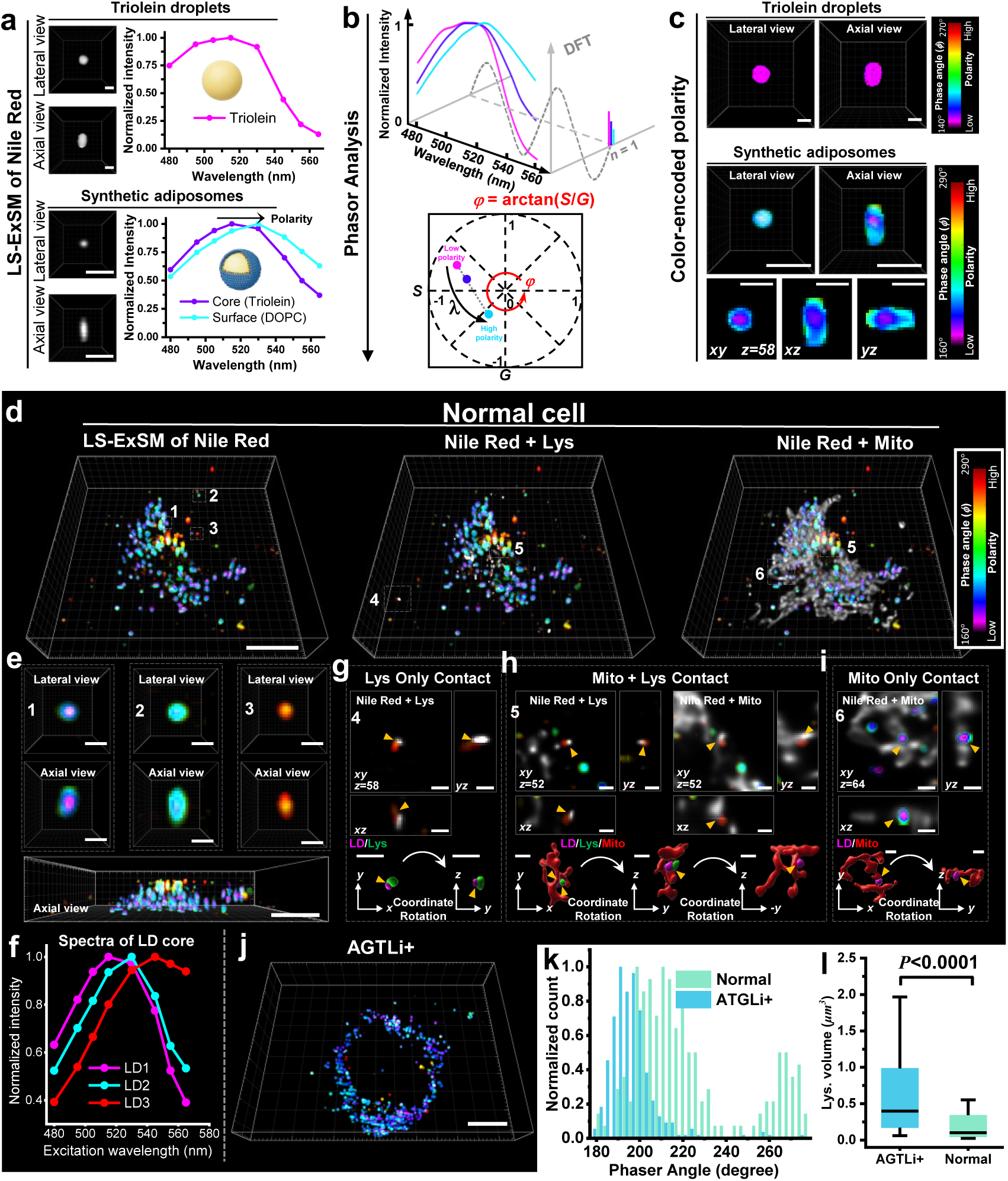
| LS-ExSM enables quantitative three-dimensional spectroscopic mapping of LD polarity and heterogeneity in cells. (a) Three-dimensional grayscale images of pure triolein droplets and adiposomes, shown in lateral and axial views. Corresponding excitation spectra (right) show uniform spectral profiles for triolein and distinct core and surface spectra for adiposomes. (b) Schematic of spectral phasor analysis. Excitation spectra are mapped to phasor coordinates (*G*, *S*) using the first harmonic of the discrete Fourier transform, with each spectrum represented as a single point (see Method). The phase angle (*φ* = arctan(*S*/*G*)) serves as a quantitative descriptor of the spectral profile, with shifts in *φ* reflecting changes in lipid polarity. (c) Three-dimensional polarity (*φ*) maps derived from spectral phasor analysis. Triolein droplets exhibit uniform polarity, whereas adiposomes show a pronounced core-to-surface gradient. Orthogonal views and representative cross-sections are shown. Color bars indicate the phase-angle ranges. (d) Volumetric polarity mapping of LDs in a COS-7 cell, with lysosome and mitochondria imaging under normal conditions. From left to right: three-dimensional LD polarity map, merged view with lysosomes (ATTO532), and merged view with mitochondria (CF568). (e) Representative LDs from (d), showing distinct internal polarity distributions. Orthogonal sections reveal polarity variation along the axial dimension. Bottom panels show the corresponding axial views from (d). (f) Excitation spectra from the three LDs with distinct polarity in (e). (g) Representative LD in contact with lysosomes. Orthogonal xy, xz and yz sections and corresponding three-dimensional rendering show elevated polarity at the contact region. (h) Representative LD simultaneously contacting lysosomes and mitochondria, shown in orthogonal sections and three-dimensional rendering. (i) Representative LD contacting mitochondria only, shown in orthogonal sections and three-dimensional rendering. (j) Volumetric LD polarity map under ATGLi+ conditions. (k) Distribution of LD phase angles (polarity) under normal and ATGLi+ conditions. (l) Box-and-whisker plots of lysosome volume under normal and ATGLi+ conditions. The center line indicates the median, and whiskers represent the 10th–90th percentiles. n = 3 cells. Scale bars, 10 μm (d, e bottom, j), 2 μm (a, c), 1 μm (e top, g, h, i).

In live COS-7 cells, LDs labeled with low concentrations of Nile Red exhibited substantial polarity heterogeneity across the whole cell (Fig. 5d and Supplementary Video 8). Representative excitation spectra from distinct LDs highlighted this variation (Fig. 5e-f), reflecting subtle but resolvable spectral shifts consistent with previous two-dimensional measurements. Importantly, the volumetric capability of LS-ExSM revealed the internal polarity organization and heterogeneity of individual LDs (Fig. 5e). Unlike two-dimensional projections, in which signals from the LD core and surface were superimposed along the axial direction, volumetric sectioning enabled depth-resolved measurements and more accurate estimation of core polarity. This resolved a key ambiguity in prior measurements, in which highly polar signals could not be distinguished from the LD surface or from surrounding membrane-associated structures. Elevated polarity was observed not only at the LD periphery but also within the droplet core (for example, droplet 3 in Fig. 5e), confirming that the high polarity was intrinsic to LDs rather than arising from surrounding membranes (Supplementary Video 9).

Correlating polarity maps with organelle localization revealed a strong association between LD polarity and lysosome interactions. LDs in contact with lysosomes—either alone or together with mitochondria—consistently exhibited higher polarity, whereas LDs contacting mitochondria alone largely retained lower polarity states (Fig. 5g-i). These relationships were consistently observed across orthogonal sections and three-dimensional renderings, demonstrating the ability of LS-ExSM to link spatial organization with local physicochemical states. Under ATGL inhibition, LD polarity shifted globally toward lower values (Fig. 5j–k). In parallel, lysosomes exhibited a marked increase in volume (Fig. 5l), which could be directly quantified in three dimensions. Despite this expansion, LD polarity remained reduced, indicating that increased lysosomal volume alone did not restore LD polarity when neutral lipolysis was inhibited. These results establish LS-ExSM as a volumetric spectroscopic imaging framework for quantitative mapping of intracellular physicochemical parameters, enabling direct measurement of functional states with subcellular spatial resolution.

## Discussion

Here we establish LS-ExSM as an approach for linking three-dimensional cellular organization to functional state through volumetric spectroscopic imaging. By encoding spectral information in the excitation domain, LS-ExSM decouples spectral acquisition from detection and captures full spectra with ∼10 nm spectral resolution throughout the imaging volume. Coupling sparse volumetric sampling with deep-learning-based reconstruction further reduces acquisition time by threefold while preserving both structural and spectral information, extending volumetric spectroscopic imaging into a high-speed regime. In this configuration, excitation spectra report not only molecular identity but also local physicochemical properties, moving fluorescence microscopy beyond intensity-based contrast toward multidimensional functional imaging.

A central advance of LS-ExSM is that it brings together spectral resolution, multiplexing capacity and volumetric imaging speed within a single platform. Conventional light-sheet microscopy typically supports simultaneous imaging of only two or three subcellular structures. Recent spectral light-sheet approaches have extended this to six colors in live cells, but still face major trade-offs among spectral resolution, acquisition speed and signal fidelity^20, 25^. Emission-dispersion-based methods generally rely on point or line scanning and therefore require minutes per volume. Excitation-based lattice light-sheet methods are faster (∼9.2 s/volume), but sparse spectral sampling (∼40 nm spacing) limits spectral resolution and causes substantial crosstalk (∼50%). Parallel detection schemes improve speed further (0.5–1 s/volume), yet divide emission into broad spectral bins (≥30 nm), sacrificing fine spectral information. In contrast, LS-ExSM captures full excitation spectra with ∼10 nm spectral resolution and achieves six-color volumetric imaging with minimal crosstalk (∼1%) at ∼0.83 volumes s⁻¹ (∼1.2 s/volume). Collecting emission within a fixed passband further improves photon efficiency and signal-to-noise ratio while avoiding dependence on the spectral response of the detection system. This combination of spectral fidelity, multiplexing and volumetric speed is difficult to achieve with existing spectral imaging approaches.

High-fidelity spectroscopic imaging in three dimensions creates new opportunities for biological interrogation. LS-ExSM enables multiplexed imaging of dynamic organelle interactions in live cells, allowing coordinated behaviors of multiple structures to be followed simultaneously with minimal crosstalk. Quantitative analysis of organelle contacts further shows that metabolic perturbations reshape interaction networks by altering contact frequency, stability, and spatial organization, while underscoring that projection-based 2D measurements can substantially overestimate contacts and even misrepresent biological trends. Volumetric spectroscopy also enables direct mapping of intracellular physicochemical parameters, such as lipid-droplet polarity, with subcellular resolution. Importantly, volumetric sectioning resolves ambiguities inherent to projection-based measurements, separating signals arising within organelles from those contributed by surrounding structures. Together, these results show how volumetric spectroscopic imaging can connect spatial organization with physicochemical state in live cells.

More broadly, LS-ExSM provides a generalizable route for using high-fidelity volumetric spectroscopy to quantify molecular composition and physicochemical states in complex biological environments. Several directions should further extend its reach. Improving spatial resolution, particularly along the axial dimension, remains an important goal, and integration with structured illumination may provide a route toward super-resolved spectroscopic imaging in three dimensions. Although organelle motion faster than the volumetric acquisition rate may introduce motion-related artefacts, spectral unmixing remains robust because separation is performed at the level of individual light-sheet planes; further gains in speed should be achievable with brighter fluorophores and faster detectors. In parallel, advances in biosensor design, especially fluorophores with larger and more informative spectral responses, should expand the range of measurable physicochemical parameters across whole-cell volumes. Taken together, these directions point toward excitation-encoded volumetric spectroscopic imaging as a broadly applicable platform for quantitative, multidimensional analysis of living systems.

## Methods

### LS-ExSM optical setup

Excitation wavelengths were selected and rapidly switched using a supercontinuum laser source (SCPro-M-40, YSL Photonics) and an acousto-optic tunable filter (AOTF; 97-03151-01, Gooch & Housego) controlled by fast electronic modulation of the radiofrequency drive signals (Supplementary Fig. S1). The output beam was linearly polarized (SHP1025, Union Optic) and filtered by a short-pass filter (FESH0750, Thorlabs) before entering the AOTF. At a conjugate plane, an aperture selected the first-order diffracted beam while blocking the zero- and higher-order components. The selected beam was coupled into a single-mode fiber (P3-S405-FC-1, Thorlabs) for spatial filtering, then collimated and relayed using achromatic lenses. The excitation beam was reshaped into an elongated profile using two cylindrical lenses (CYX0007 and CYX0012, Union Optic). A mechanical slit placed at the Fourier plane limited the effective numerical aperture of the illumination, and a third cylindrical lens (CYX0009, Union Optic) focused the beam along one axis to generate a one-dimensional Gaussian profile. The beam was then reflected by a dichroic mirror (ZT561rdc, Chroma) and relayed through a 4f system to a galvo mirror (Scanlab GmbH), which was conjugated to the back focal plane of the primary objective (O1, Olympus PlanApo N 60× oil, NA = 1.42). By introducing an off-axis displacement at the pupil plane of O1, the excitation beam formed an oblique light sheet at the sample plane with a tilt angle of 30°. Scanning of the galvo mirror translated the light sheet across the sample, while the relay optics maintained pupil conjugation between the scanning mirror and O1.

Fluorescence emission was collected by O1 and propagated back through the same optical path. After passing through the dichroic mirror, the emission was relayed by a second 4f system to a secondary objective (O2, Nikon CFI TU Plan Fluor EPI 100×, NA = 0.9). The pupil planes of O1 and O2 were conjugated by relay lenses, and the galvo mirror performed descanning so that the intermediate image at the remote plane remained stationary during scanning. The intermediate image was then re-imaged by a tertiary objective (O3, Olympus LUMPlanFL N 60× water, NA = 1.00) and projected onto a scientific CMOS camera (Zyla 4.2 Plus, Andor) through achromatic lenses. A custom water chamber at O3, following the design of Yang et al.^13^, was implemented to increase the effective numerical aperture of the detection path. An emission filter (FF01-650/150-25, Semrock) was placed before the camera to reject residual excitation light. The optical configuration satisfied both the sine and Herschel conditions through pupil conjugation between O1 and O2 using two 4f relay systems. The lateral magnification from the sample plane to the intermediate image was set to 1.5, matching the refractive index of the immersion medium (n = 1.518), which also yielded an axial magnification of 1.5. The overall system magnification was 100×. The sCMOS camera has a physical pixel size of 6.5 μm; with 2 × 2 binning, the effective pixel size in the sample plane was 130 nm, satisfying Nyquist sampling.

### Architecture and implementation of the deep-learning restoration algorithm for sparse volumetric spectroscopic imaging

#### Data preprocessing

Raw fluorescence microscopy data were acquired with a 16-bit dynamic range to preserve linear intensity response and fine signal variations. Although the framework did not require strict global normalization, localized intensity rescaling was applied to facilitate stable convergence during training. To maximize data utilization, a slice-wise partitioning strategy was adopted. For denoising, volumes were decomposed into sub-stacks according to spectral channel and axial position. For intermediate slice reconstruction (ISR), training pairs were generated by subsampling axial or temporal stacks, thereby expanding a limited number of biological samples into a larger training dataset.

#### Self-supervised spatio-temporal denoising

The denoising network operated directly on noisy observations without requiring clean ground-truth references. To exploit both spatial redundancy and temporal correlation, the network used a spatio-temporal input window consisting of five adjacent frames. Self-supervised training was based on an orthogonal masking strategy. At each iteration, two sub-sampled volumes, 𝑀_𝐴_(𝑥) and 𝑀_𝐵_(𝑥), were generated by selecting neighboring pixels on complementary grids. The network 𝑓(⋅) was optimized by minimizing the discrepancy between the prediction from 𝑀_𝐴_(𝑥) and the target 𝑀_𝐵_(𝑥).

#### Intermediate slice reconstruction (ISR)

The ISR module was designed to generalize across diverse biological structures. Training pairs were generated from sparsely sampled axial or temporal stacks, yielding approximately 300 pairs from 2–3 biological samples. To preserve structural fidelity, the ISR network was trained using a composite loss function that combined intensity reconstruction with edge-aware supervision. Edge features were extracted using a gradient-based operator (for example, Sobel) to compute an edge consistency loss (ℒ_𝐸_), which penalized structural discrepancies between predicted and ground-truth slices.

#### Implementation and training protocol

All models were implemented in PyTorch and trained on a workstation equipped with an NVIDIA RTX 4070 GPU (12 GB VRAM). Optimization was performed using the AdamW optimizer with an initial learning rate of 2×10^−4^, together with a cosine annealing schedule. Models were trained for 200 epochs with a batch size of 8. A typical training run required 2–4 h.

#### Quantitative evaluation metrics

Performance was evaluated using local signal-to-noise ratio (SNR), structural similarity index (SSIM), and spectral angle mapping (SAM), which quantified denoising performance, structural fidelity, and spectral fidelity, respectively.

### Image acquisition

System timing was controlled using two data acquisition (DAQ) devices. A PCI-6733 (National Instruments) synchronized camera exposure with the AOTF radiofrequency drive signals for excitation wavelength switching. A USB-6211 (National Instruments) synchronized the camera frame signal with the galvo mirror control voltage, thereby defining the beam offset at the objective pupil and controlling light-sheet scanning across the sample. Consecutive frames corresponded to different excitation wavelengths within a predefined spectral cycle. After completion of each spectral cycle, the galvo mirror position was updated and the wavelength sequence was repeated. Eight excitation wavelengths (565, 555, 545, 530, 515, 505, 495 and 480 nm) were used for spectral acquisition, with relatively small spacing to preserve spectral fidelity. Because the galvo mirror was the only moving component, the volumetric imaging rate was determined by the camera frame rate. For six-color live-cell imaging, the camera was operated at 217 Hz with a region of interest (ROI) of 1056 × 320 pixels to balance signal level, photobleaching, and phototoxicity. A total of 33 planes were acquired at a plane interval (z′) of 1.12 μm by sparse volumetric sampling and subsequently reconstructed to 97 planes using the deep-learning framework, yielding a field of view of 90 × 70 × 10 μm³. In total, 198 volumes were acquired continuously. The excitation power at each wavelength before entering O1 was 33 μW. For high-speed volumetric imaging of the ER, the frame rate was increased to 938 Hz using a reduced ROI of 1056 × 152 pixels. A total of 33 planes were acquired by sparse volumetric sampling and subsequently reconstructed to 97 planes using the deep-learning framework, corresponding to a field of view of 90 × 70 × 5 μm³. Imaging was performed at 545 nm with an excitation power of 90 μW before entering O1, and no obvious photobleaching of the stable fluorescent protein mGold was observed over 300 volumes^40^.

### Multi-target image processing and organelle interaction analysis

Raw image stacks from six-color live-cell imaging and single-channel high-speed volumetric imaging were first processed using a self-supervised denoising network to improve the signal-to-noise ratio. The denoised data were then input into a supervised reconstruction network with edge-aware constraints to infer intermediate slices and recover complete volumetric datasets from sparsely sampled inputs. The reconstructed data were linearly unmixed into individual fluorophore channels using a custom MATLAB implementation. Non-negative least-squares regression was applied, in which the excitation spectrum at each pixel was expressed as a linear combination of reference spectra. Each unmixed channel was subsequently deconvolved in Huygens Professional (Scientific Volume Imaging) using experimentally measured point spread functions. The data remained in the native skewed coordinate system (x′–y–z′) and were processed using a deskewing algorithm implemented in the PetaKit5D toolbox^41^, followed by rotation to recover the standard Cartesian (x–y–z) coordinate system. Organelle segmentation and contact analysis were performed in Imaris (Bitplane). Surfaces were generated for each channel using manually adjusted intensity thresholds. Touching objects were separated using the “split touching objects” function for approximately spherical organelles, including LDs, peroxisomes and lysosomes. Objects smaller than 10 voxels were excluded from further analysis.

### Spectral phasor analysis of LD polarity

For Nile Red–labeled lipid droplets, excitation spectral phasor analysis was performed on a per-voxel basis. The excitation spectrum at each voxel was mapped to phasor coordinates (*G*, *S*) using the first harmonic of the discrete Fourier transform.

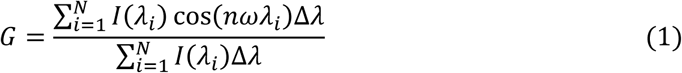

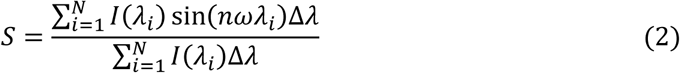

𝜆_𝑖_ represents the excitation wavelength of the *i*-*th* spectral channel; 𝐼(𝜆_𝑖_) is the collected fluorescent intensity of the *i*-*th* channel; 𝜔 = 2𝜋/𝑁 with *N* is the number of spectral channels (here, *N* = 8), and *n* is the harmonic order (here, *n* = 1). The phase angle is expressed as,

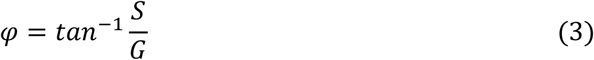

The phase angle *φ* is directly related to the center of mass of the spectrum, and used as a quantitative descriptor of the spectral profile. Phase-angle maps were generated and visualized as color-coded volumetric renderings. The resulting data were then deskewed and rotated to transform from the native (x′–y–z′) coordinate system to the standard (x–y–z) coordinate system.

### Cell culture and fixation

COS-7 cells (Cell Resource Center, IBMS, CAMS/PUMS) were plated on 18 mm glass coverslips in 12-well plates and cultured in Dulbecco’s modified Eagle’s medium (DMEM; C11995500BT, Gibco) supplemented with 10% fetal bovine serum (Gemini) and 1× non-essential amino acids (11140050, Gibco). Cells were maintained at 37 °C in 5% CO₂. After 24 h, cells were fixed with 3% paraformaldehyde (E672002, Sangon Biotech) and 0.1% glutaraldehyde (16020, Electron Microscopy Sciences) in phosphate-buffered saline (PBS). Residual aldehydes were quenched by two washes with 0.1% sodium borohydride in PBS, followed by three washes in PBS (10 min each).

### Plasmid constructs and transfection

tdTomato-ER-3 and mOrange2-Peroxisomes-2 were obtained from Michael Davidson (Addgene #58097 and #54596). Mito-PhiYFP (pPhi-Yellow-mito) was obtained from Evrogen (#FP607). LAMP1-mGFP was a gift from Esteban Dell’Angelica (Addgene #34831). LAMP1-tdTomato was generated by replacing EGFP in LAMP1-mGFP with tdTomato using BamHI and NotI restriction sites. mGold-ER-3 was constructed by replacing tdTomato in tdTomato-ER-3 with a synthesized mGold sequence (Addgene #158007) using AgeI and BamHI, followed by addition of a C-terminal KDEL sequence for ER retention. Transient transfections were performed using Lipofectamine 2000 (11668019, Thermo Fisher Scientific) according to the manufacturer’s instructions.

### Multi-target labeling of live-cell samples

For six-structure labeling, cells were transiently transfected with mOrange2-Peroxisomes-2, Mito-PhiYFP and mGold-ER-3. Prior to imaging, cells were incubated with LysoTracker Red (3.5 μM, C1046, Beyotime) for 1 h, SYBR Gold (1:40,000, S11494, Thermo Fisher Scientific) for 20 min and LipidSpot488 (LS488; 1:1500, 70065, Biotium) for 5 min. Cells were then washed three times with DPBS (5 min each). For four-structure labeling, cells were transfected with Mito-PhiYFP, LAMP1-tdTomato and mGold-ER-3, followed by LS488 staining. For three-structure labeling, cells were transfected with Mito-PhiYFP and LAMP1-tdTomato, followed by LS488 staining. For inhibition of neutral lipolysis, cells were treated with 25 nM atglistatin (ATGL inhibitor; 15284, Cayman Chemical) in complete medium for 24 h at 37 °C in 5% CO₂ after transfection.

### Multi-target labeling of fixed-cell samples

Cells were permeabilized and blocked in buffer containing 3% bovine serum albumin (BSA) and 0.1% Triton X-100 (Sigma) in PBS, followed by primary and secondary antibody staining. Primary antibodies included rat anti-α-tubulin (MAB1864, Millipore) for microtubules, mouse IgG1 anti-GM130 (610822, BD Transduction Laboratories) for the Golgi apparatus and mouse IgG2a anti-Tom20 (sc-17764, Santa Cruz Biotechnology) for mitochondria. The following dye-conjugated secondary antibodies were prepared via reaction with dye NHS esters: goat anti-rat IgG (H + L) (Jackson ImmunoResearch 112-005-167)-CF514 (92103, Biotium), goat anti-mouse IgG1 (Jackson 115-005-205)-Atto542 (AD542-31, ATTO-TEC), goat anti-mouse IgG2a (Jackson 115-005-206)-CF568 (92131, Biotium). After primary antibody labeling and prior to secondary antibody staining, cells were incubated with SYBR Gold (1:400,000) in washing buffer (0.3% BSA, 0.01% Triton X-100 in PBS) for 5 min to label nuclear nucleic acids. Following antibody staining, cells were incubated with YF633-phalloidin (2 T/mL, YP0053, UElandy) for 20 min to label actin. Finally, cells were stained with LS488 (1:1500) for 5 min to label lipid droplets and washed three times with PBS (10 min each) before imaging.

### LS-ExSM of Synthetic Lipid Droplets in vitro

A stock solution of triolein (TAG; HY-N1981, MedChemExpress) was prepared at 100 mg mL⁻¹ in hexane. 1,2-Dioleoyl-sn-glycero-3-phosphocholine (DOPC; HY-113424A, MedChemExpress) was prepared at 50 mg mL⁻¹ in a 1:1 (v/v) mixture of hexane and chloroform. Triolein droplets were generated by mixing 8 μL of lipid stock with 290 μL deionized water (diH₂O) and 1–2 μL Triton X-100, followed by vortexing and sonication for 2–3 min. Artificial adiposomes were prepared by mixing triolein and DOPC at a 2:1 molar ratio. The solvent was evaporated under a nitrogen stream, followed by addition of 500 μL diH₂O. The suspension was heated in a boiling water bath to melt the lipids, immediately sonicated for 30 min, diluted, stained with 1 μM Nile Red, and mounted on 2% BSA-coated coverslips prior to imaging.

### LS-ExSM of LDs in cells with lysosome and mitochondria labeling

Live cells were incubated with Nile Red (30 nM) for 10 min to label lipid droplets. After staining, cells were washed with DPBS for 5 min and imaged by LS-ExSM. Following live-cell imaging, cells were fixed on the microscope stage using ethanol for 10 min to remove Nile Red signal, and washed three times with PBS (10 min each). Cells were then permeabilized and blocked in blocking buffer for 1 h at room temperature. Primary antibodies against LAMP1 (mouse anti-LAMP1; 555798, BD Transduction Laboratories) and Tom20 were applied simultaneously in blocking buffer for 1 h, followed by three washes in washing buffer (10 min each). Cells were then incubated with goat anti-mouse IgG1–ATTO532 and goat anti-mouse IgG2a–CF568 secondary antibodies in blocking buffer for 30–60 min at room temperature. Finally, samples were washed three times with washing buffer (10 min each) and once with PBS (10 min) prior to imaging.

## Acknowledgments

This work was supported by the National Natural Science Foundation of China (62475032, 62205048), the Sichuan Science and Technology Program (2026YFHZ0044) and the National Research Foundation of Korea (NRF) grants funded by the Korean government (RS-2024-00339900).

## Author Contributions

K.C. conceived the research. J.Y. designed and conducted the experiments. J.X. developed the neural network framework for sparse volumetric imaging. All authors contributed to experimental designs, data analysis, and paper writing.

